# Serine/threonine protein kinase phosphorylation of DosR alters target gene transcription mechanics and regulates *Mycobacterium tuberculosis* sensitivity to nitric oxide stress

**DOI:** 10.1101/2025.09.03.674086

**Authors:** Natalie R. Sontag, Ana Ruiz Manzano, Alwyn M. V. Ecker, Eric A. Galburt, Shumin Tan

## Abstract

Successful host colonization by bacterial pathogens requires appropriate response and adaptation to environmental signals encountered during infection, with two-component systems (TCSs) and serine/threonine protein kinases (STPKs) being two important signal transduction mechanisms. *Mycobacterium tuberculosis* (Mtb) possesses similar numbers of STPKs (11) and TCSs (12), but if and how these two regulatory systems coordinate to enable Mtb adaptation in response to key environmental cues remains poorly understood. Here, we identify extensive interactions between STPKs and TCSs, with a subset of STPKs demonstrating interactions with multiple TCS response regulators. STPK phosphorylation of DosR, the response regulator of the key nitric oxide (NO)/hypoxia-responsive TCS DosRS(T), decreased its binding to target promoter DNA and its ability to activate steady-state gene transcription, in marked contrast with the opposite phenotypes observed with the activated, phospho-aspartic acid form of DosR. Strikingly, a ΔSTPK Mtb mutant exhibited increased DosR regulon transcription at lower NO levels than wild type Mtb, illustrating how STPK phosphorylation of a TCS RR may act to restrict and fine-tune conditions in which activation occurs. Together, our results support a functional relationship between STPKs and TCSs, and shed light on the mechanisms underpinning STPK-TCS interplay.

**AUTHOR SUMMARY:** *Mycobacterium tuberculosis* (Mtb) is the bacterium that causes tuberculosis, which remains the largest cause of death from an infectious disease globally. Successful host colonization by Mtb requires that the bacteria appropriately sense and respond to changes encountered in its local microenvironment throughout the course of infection. Here, we provide evidence for the interplay between two key signal transduction regulatory mechanisms – two-component systems (TCSs) and serine/threonine protein kinases (STPKs). Focusing on the DosRS(T) TCS that is crucial in the response of Mtb to the critical environmental signals of nitric oxide (NO) and hypoxia, we reveal that STPK phosphorylation of the DosR regulator decreases target gene promoter binding and the activation of steady-state transcription. Further, an Mtb mutant that was disrupted in an STPK that phosphorylates DosR exhibited increased DosR target gene expression at lower NO concentrations than wild type Mtb. These results indicate that STPK phosphorylation serves as an additional regulatory layer for TCSs, adjusting the DosR concentration range under which full activation of the TCS occurs.

## INTRODUCTION

The ability of *Mycobacterium tuberculosis* (Mtb) to sense and respond to dynamic changes in environmental signals encountered throughout the course of infection is critical for successful host colonization. This includes cues such as acidic pH, chloride (Cl^-^), potassium (K^+^), nitric oxide (NO), and hypoxia, which are associated with the host immune response [1–7]. Mtb is the causative agent of tuberculosis and is the leading global cause of death from an infectious disease [8]. The heterogeneity of the environment Mtb experiences during host infection further affects the bacterial physiological state and contributes importantly to the difficulty of tuberculosis treatment [9–13].

Phosphorylation-based signal transduction enables environmental adaptation by linking extracellular signals to intracellular regulatory mechanisms. In bacteria, the best-studied mechanism of phosphorylation-based transmembrane signaling are two-component systems (TCSs), where ligand binding by the transmembrane histidine kinase (HK) sensor protein initiates a phosphorelay to the cognate intracellular response regulator (RR) protein, typically a transcription factor that controls expression of specific genes [14, 15]. The number of TCSs per bacterial genome strongly correlates with ecological and environmental niche [16, 17] – bacteria living in more constant environments usually encode fewer TCSs, while bacteria that inhabit rapidly changing or diverse environments typically encode larger numbers of these signaling proteins that are critical for cellular processes [18–20]. Mtb encodes 12 TCSs that play a key role in virulence, environmental adaptation, and infection [14]. For example, inactivation of PhoPR, a key TCS involved in Mtb response to acidic pH and Cl^-^ [3, 5], results in significant attenuation in the ability of Mtb to colonize its host [21]. An essential TCS in Mtb, PrrAB, was reported to be involved in early adaptation to intracellular infection [22, 23], and we have since shown that PrrA is a global regulator of Mtb response to acidic pH, high [Cl^-^], hypoxia and NO [24]. The TCS DosRS(T) regulates Mtb response to hypoxia and NO, and upon sensing of these signals, mediates induction of a “dormant” state through the control of 50 genes known as the “dormancy regulon” [4, 6, 7, 25]. The TCS KdpDE is responsible for regulation of the Kdp K^+^ uptake system and is pivotal for Mtb adaptation to low environmental potassium levels ([K^+^]) [1, 26]. These examples illustrate the importance of TCSs for Mtb adaptation to its local environment.

Importantly, in addition to TCSs, signal transduction and environmental response in Mtb is also mediated by serine/threonine protein kinases (STPKs) [27–29]. In contrast to TCSs, STPKs have a larger set of phosphorylation targets, and while STPKs are often less numerous in the genome of a given bacterium, their widespread presence and impact on bacterial biology have become increasingly appreciated [27–32]. For example, phosphorylation of glutamyl tRNA reductase by the STPK Stk1 plays an important role in the regulation of heme biosynthesis in *Staphylococcus aureus* [33]. Another example is with *Listeria monocytogenes*, where phosphorylation of the protein ReoM by the STPK PrkA is essential for viability, due to its role in peptidoglycan synthesis [34]. The Mtb genome markedly contains a comparatively large number of STPKs compared to other bacterial species, with 11 STPKs encoded [27, 30, 35]. Notably, global phosphoproteome studies have identified TCS RRs as potential substrates of STPK phosphorylation [31, 32], raising the concept of STPK-TCS interplay in the regulation of TCS function. Indeed, there is increasing support for this interplay, with studies in bacterial species ranging from Mtb and *Bordetella*, to *Streptococcus pneumoniae*, *S. aureus*, and *Bacillus subtilis* [31, 32, 36–42]. Specifically for Mtb, all 12 TCS RRs have been identified from global phosphoproteome studies as potential substrates of STPK phosphorylation [31, 32]. In the case of DosR and PrrA, both RRs from TCSs that respond to NO stress among other signals, biochemical assays with purified proteins have further verified their phosphorylation by STPKs [36, 43], with the STPK PknH shown to phosphorylate DosR [36]. Strikingly, we have shown that a STPK phosphoablative PrrA-T6A Mtb mutant was significantly altered in its transcriptional response to acidic pH and high [Cl^-^], and dampened in environmental response to both NO and hypoxia [24]. Consequently, this mutant was incapable of entry into an adaptive state of growth arrest upon extended exposure to NO and attenuated for host colonization *in vivo* [24]. While there is clear evidence for interplay between STPKs and TCSs, much remains unknown regarding how the two systems interact in the coordination of Mtb environmental adaptation.

Here, we examined STPK interactions with the RRs DosR, PrrA, PhoP, and KdpE, all RRs known to play critical roles in Mtb transcriptional response to environmental cues encountered throughout host infection [1, 3–7], revealing both specificity and overlap in interactions. Focusing on DosR as an exemplar system, we find that PknH and PknD phosphorylation of DosR decreased its binding to the promoter of its target genes, with strong binding restored by combined treatment with acetyl phosphate (AcP), which mimics HK phosphotransfer [44, 45]. Further, PknH and PknD phosphorylation of DosR decreased the steady-state transcription rate of DosR target genes, in contrast to the increase observed upon AcP treatment of DosR. Additionally, STPK phosphorylation shifted the AcP-treated DosR concentration dependence of target gene activation. Finally, we found that a Δ*pknH* Mtb mutant exhibited increased DosR regulon transcription at lower NO levels than wild type (WT) Mtb. Combined, our work sheds light on the mechanisms underpinning STPK-TCS interplay, illustrating how STPK phosphorylation of a TCS RR can act to restrain its activation to ensure response initiation only when appropriate.

## RESULTS

### There is both overlap and specificity in interactions between STPKs and TCS RRs

To systematically identify possible interactions between STPKs and TCSs RRs, we utilized the mycobacterial protein fragment complementation (M-PFC) assay [46]. Comparable to the bacterial two-hybrid assay, this method is based on functional reconstitution of the murine dihydrofolate reductase (mDHFR) driven by interactions between two test proteins, thereby conferring resistance to trimethoprim (TRIM) [46]. A previous study applying this method uncovered an interaction between DosR and the STPK PknH, providing precedence for its use in this context [36]. In particular, we examined the RRs DosR, PrrA, PhoP, and KdpE, fusing the open reading frame of each to the mDHFR fragment F1,2, with each of 9 STPKs fused to the mDHFR fragment F3. Some combinations of RRs and specific STPKs showed clear growth at 50 µg/mL TRIM, indicating a strong interaction (Figs 1 and 2). As controls, *Mycobacterium smegmatis* containing empty vectors were unable to grow on 7H10 TRIM plates, whereas all strains grew well on 7H10 plates lacking TRIM (Fig 1). We compared interactions across four RRs to explore possible relationships between environmental signals responded to by a TCS and interactions with STPKs, as shown in Fig 2. Interestingly, we observed almost complete overlap in the STPKs that interact with PrrA and DosR, with smaller subsets of those same STPKs interacting with PhoP and KdpE. These results indicate that there is both overlap and specificity in the interactions between STPKs and RRs.

**Fig 1.**
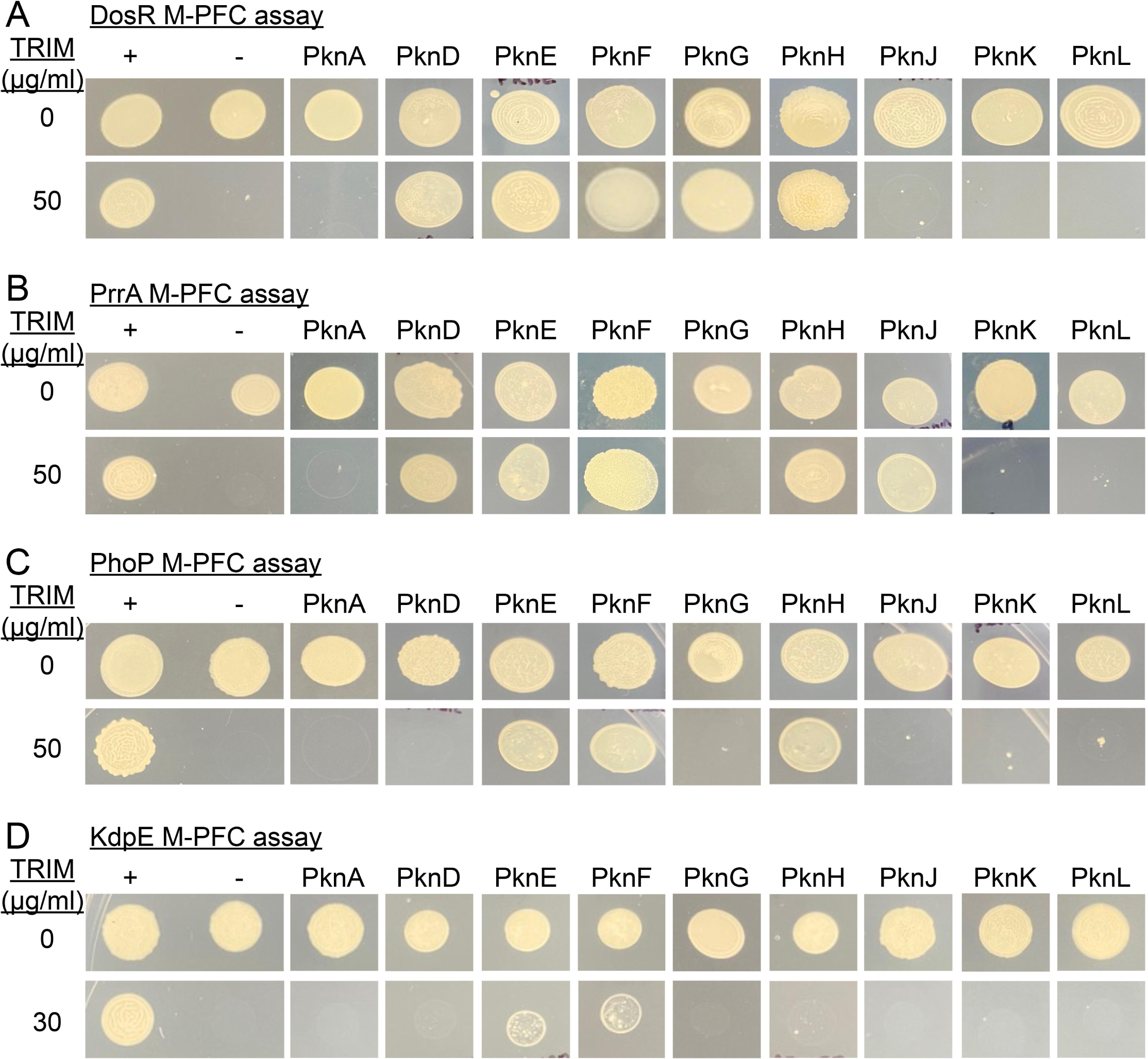
Specificity of interactions between TCS Res and STPKs. Interactions between the TCS RRs DosR (A), PrrA (B), PhoP (C), or KdpE (D) with the various STPKs (kinase domains only for all except PknG and PknK) were tested by M-PFC assay. The positive control (“+”) was *M. smegmatis* expressing the *S. cerevisiae* GCN4 dimerization domains fused to the F1,2 or F3 domains of the murine dihydrofolate reductase gene; the negative control (“-“) was *M. smegmatis* expressing the respective RR fused to the F1,2 domain and a F3 domain that was not fused to any Mtb gene. Data are representative of 3 independent experiments.

**Fig 2.**
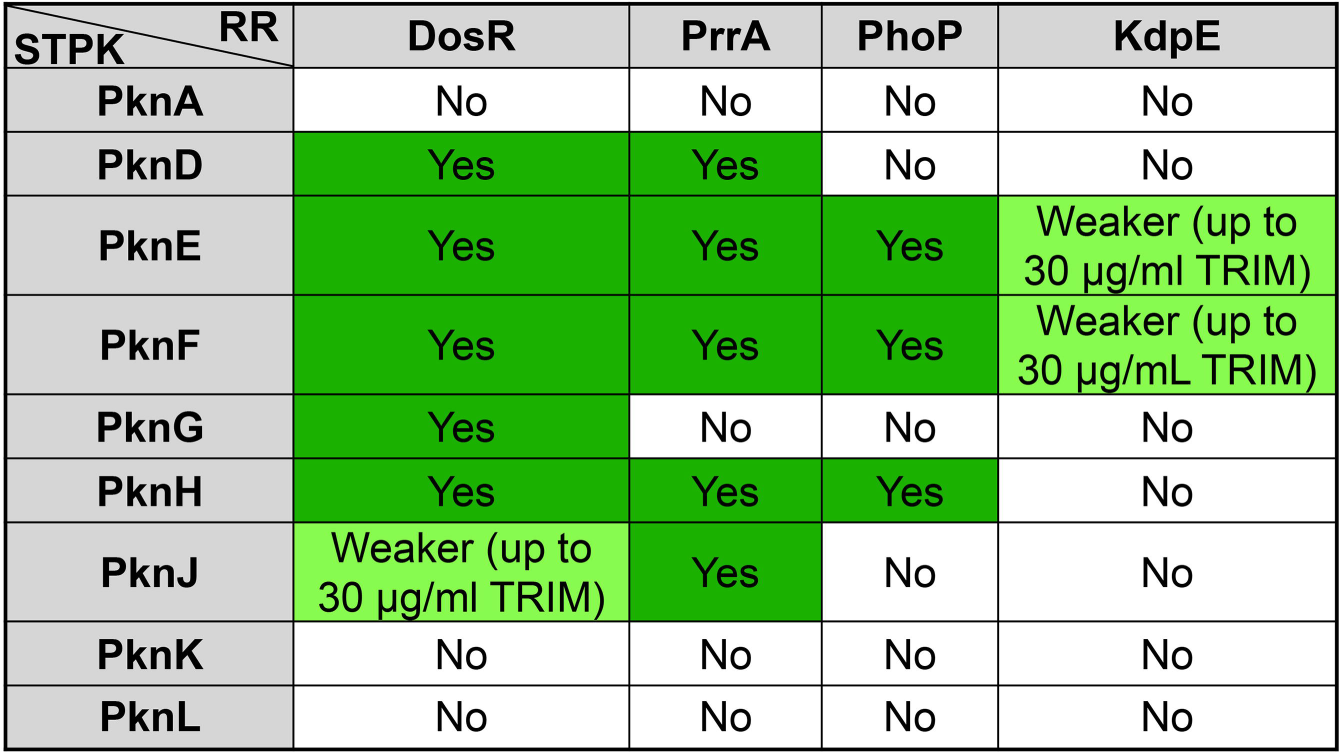
There is both overlap and specificity in interactions between STPKs and TCS RRs. Summary table of interactions between the RRs PrrA, DosR, PhoP, and KdpE with various STPKs (kinase domains only for all except PknG and PknK) as determined by M-PFC assays. Results are representative of 2-3 independent experiments.

In accord with previous results [36], mass spectrometry (MS) analysis of recombinantly expressed and purified DosR that was subsequently phosphorylated *in vitro* with recombinant PknH identified phosphorylation at the Thr198 and Thr205 residues. Analysis of DosR samples also directly confirmed that AcP treatment, mimicking HK phosphotransfer [44, 45], led to phosphorylation of the Asp54 residue [47, 48]. Notably, analyzing the results from a previous global study of the Mtb O-phosphoproteome, where phosphosites in individual STPK overexpression and deletion mutants were compared to WT Mtb [32], provided support for the likely physiological interactions of several STPK-TCS RRs indicated by our M-PFC assay results. In particular, significant differences in phosphopeptides within DosR (*i.e.*, increased presence upon STPK overexpression and/or decreased presence with the STPK deletion mutant, as compared to WT Mtb) were identified for all of the STPKs identified as DosR interactors in our M-PFC assay [32]. This was similarly the case for PhoP, while significant differences in phosphopeptides within PrrA were identified for PknD and PknE in the global phosphoproteome study [32]. Together, our results reinforce the concept of extensive interplay between STPKs and TCS RRs in Mtb.

### Changes in DosR phosphorylation status alter its binding affinity to target promoters

TCS RRs are commonly transcription factors that mediate their activity through changes in gene transcription [14, 15]. To examine how STPK phosphorylation of TCS RRs mechanistically affect their function, we thus first analyzed effects on target promoter binding, focusing our studies on DosR as an exemplar RR. As expected, AcP treatment to mimic HK phosphotransfer enhanced DosR binding to the promoter of *hspX*, a member of the DosR regulon [7, 25, 45], as indicated by electrophoretic mobility shifts observed at lower DosR concentrations with AcP-treated DosR (Figs 3A and 3B). Intriguingly, *in vitro* phosphorylation of DosR with the STPK PknH or PknD resulted in decreased DNA binding affinity to a similar degree (compare Figs 3C and 3D to Fig 3A). In a live bacterium, both HK phosphotransfer and STPK phosphorylation might be expected to co-occur at times, therefore we next examined the influence of both types of phosphorylation on DosR target promoter binding affinity. AcP treatment of PknH or PknD phosphorylated DosR resulted in a DNA binding affinity that resembled that of AcP-treated DosR alone (compare Figs 3E and 3F to Fig 3B). Similar changes in promoter binding affinity depending on DosR phosphorylation status were observed with DosR binding to the promoter of *fdxA*, another member of the DosR regulon (S1 Fig) [7, 25, 49]. Complementary to the electrophoretic mobility shift assays, we utilized fluorescence polarization to quantitatively measure DosR binding to target gene promoters. Consistent with the EMSA results, the dissociation constant (K_d_) of DosR binding to its target *hspX* and *fdxA* promoters was significantly increased upon PknH or PknD phosphorylation of DosR, demonstrating a decrease in binding affinity (Fig 4). Together, these results show that STPK phosphorylation of DosR alone, in the absence of HK phosphotransfer, decreases the affinity of DosR for its target gene promoters, indicating that STPK phosphorylation can serve as a modulatory mechanism providing tighter control of DosR activation.

**Fig 3.**
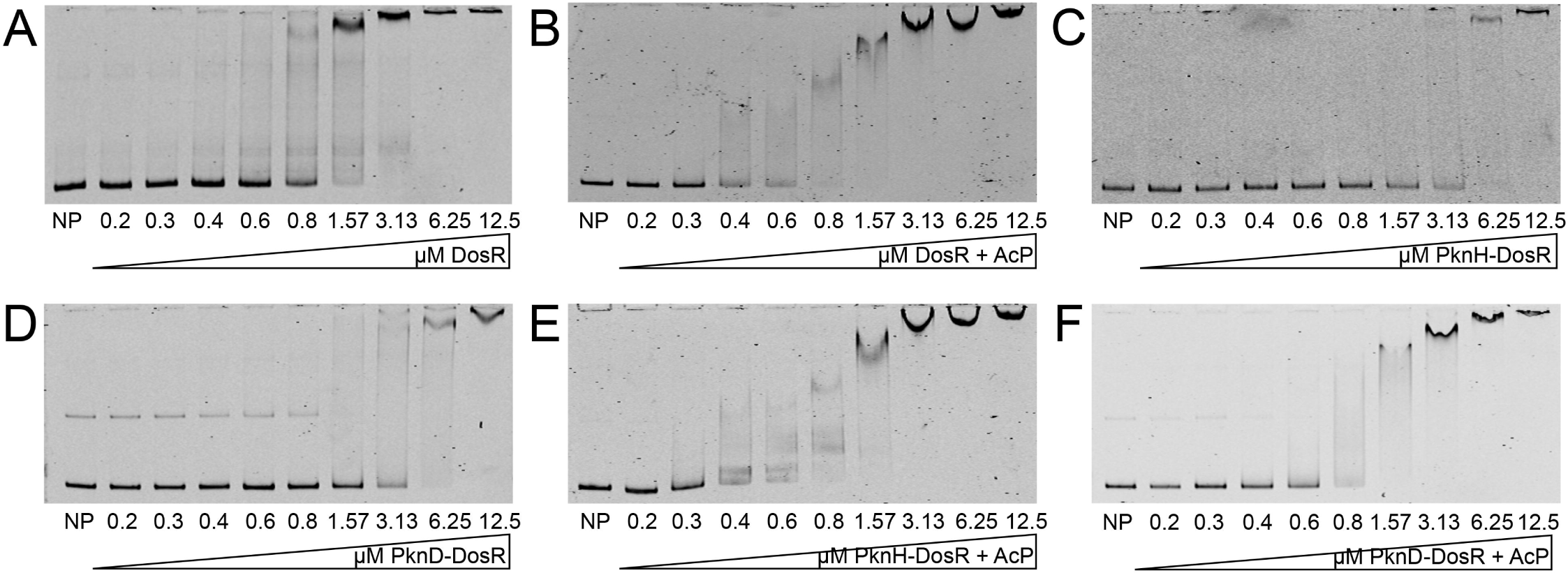
Changes in DosR phosphorylation status alter binding affinity to the promoter of its target gene *hspX*. Electrophoretic mobility shift assays (EMSAs) using purified recombinant C-terminally 6x-His-tagged DosR and IRDye 700-labeled probes for the *hspX* promoter are shown. A control with no protein (“NP”) added is shown for each gel. DosR was added at indicated concentrations for all other lanes. 40 fmoles of *hspX* promoter DNA was used in each reaction. EMSAs shown are as follows: (A) untreated DosR, (B) DosR incubated with 50 mM acetyl phosphate (AcP), (C) DosR phosphorylated “on-bead” with 1 µM PknH, (D) DosR phosphorylated “on-bead” with 1 µM PknD, (E) DosR phosphorylated “on-bead” with 1 µM PknH, then purified and incubated with 50 mM AcP, and (F) DosR phosphorylated “on-bead” with 1 µM PknD, then purified and incubated with 50 mM AcP. Data are representative of 3 independent experiments.

**Fig 4.**
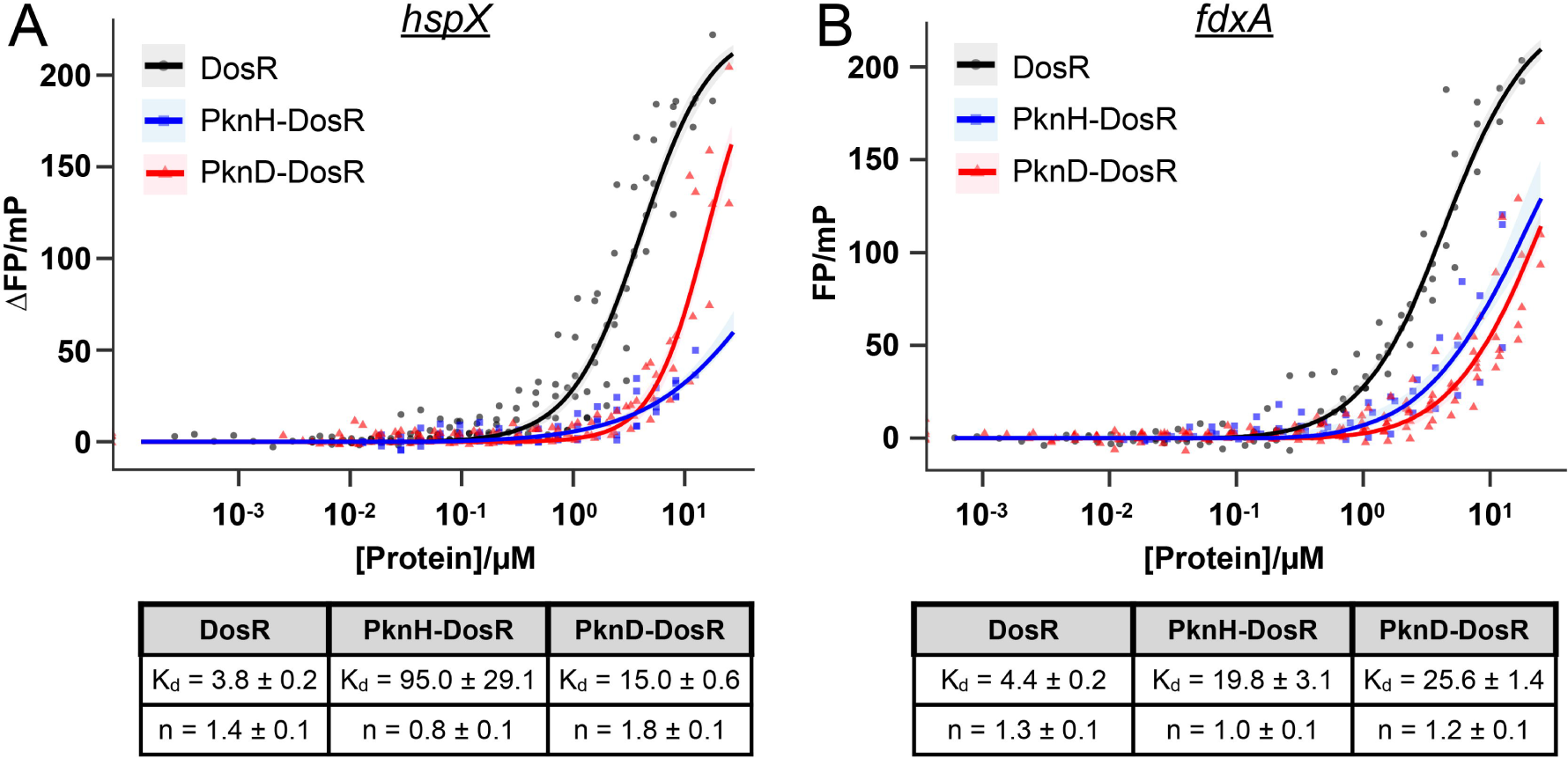
Fluorescence polarization assays demonstrate inhibition of DosR binding to its target gene promoters upon PknH or PknD phosphorylation. The change in fluorescence anisotropy (ΔFP) relative to no protein was measured as a function of increasing concentrations of DosR (black curves), PknH-phosphorylated DosR (blue curves), or PknD-phosphorylated DosR (red curves) incubated with a fluorescently labeled *hspX* (A) or *fdxA* (B) promoter DNA region. The fit using a Hill equation to estimate binding parameters with 95% confidence intervals (shaded regions) are shown. The dissociation constants (Kd) and Hill coefficients (n) are indicated in the tables below the graphs.

### STPK phosphorylation of DosR decreases steady-state transcription rates of its target genes

While EMSAs and fluorescence polarization provide insight into target promoter DNA binding, RR promoter binding affinity is just one factor determining its effect on gene transcription. For example, a transcription factor with high DNA-binding affinity but with reduced ability to interact productively with RNA polymerase [50, 51], or to modulate the kinetics of transcription initiation [52], would not be expected to be a strong transcriptional activator. To investigate the effects of STPK phosphorylation on RNA production directly, we utilized a fluorescent RNA aptamer-based method to quantify the effect of STPK phosphorylation of DosR on the steady-state rate of target gene transcription. This assay exploits the use of a Spinach-mini aptamer that produces a fluorescence-based enhancement when binding a small molecule fluorophore, such that each transcription event results in a consequent increase in fluorescence, allowing for transcript production to be monitored in real time [53, 54]. Examining *fdxA* as the target gene, phosphorylation of DosR with PknH or PknD inhibited the ability of DosR to increase steady-state transcription (Fig 5A), corresponding with the decrease in binding to the *fdxA* promoter observed with PknH or PknD-phosphorylated DosR (Figs 3C, 3D, and 4).

**Fig 5.**
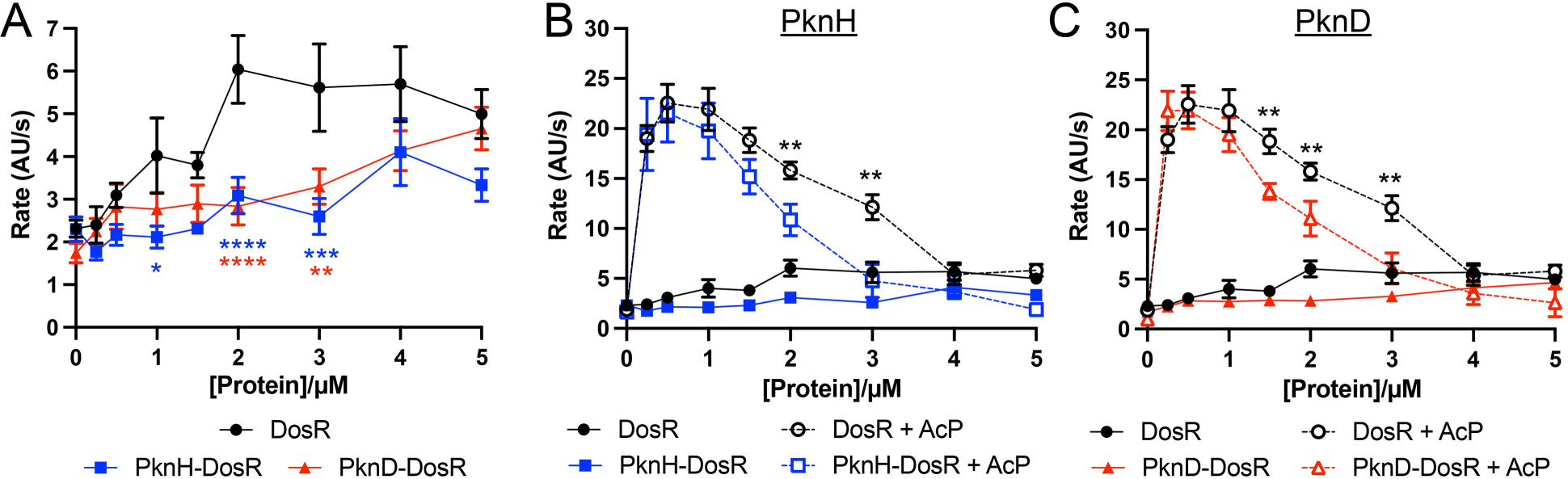
STPK phosphorylation of DosR decreases steady-state transcription dynamics of target genes. (A) STPK phosphorylation of DosR inhibits the ability of DosR to increase target gene steady-state transcription. A Spinach RNA aptamer assay was run with the *fdxA* promoter with different concentrations of indicated DosR protein. For “PknH-DosR” and “PknD-DosR”, phosphorylation of DosR with PknH or PknD, respectively, was performed “on-bead” before final purification of the phosphorylated DosR utilized in the assay. Fluorescence (arbitrary units, “A.U.”) was tracked over time on a plate reader, and steady-state rate calculated. Data are shown as means ± SEM from 3-8 experiments. p-values were obtained with a 2-way ANOVA with Tukey’s multiple comparisons. p-value in blue and red correspond to those for PknH-DosR and PknD-DosR, respectively, as compared to DosR. * p<0.05, ** p<0.01, *** p <0.001, **** p<0.0001. (B and C) STPK phosphorylation alters the dynamics of activated DosR. A Spinach RNA aptamer assay was run with the *fdxA* promoter with different concentrations of indicated DosR protein, as in (A). 50 mM acetyl phosphate (AcP) was used where indicated. Data are shown as means ± SEM from 3-8 experiments. WT, PknH-DosR, and PknD-DosR data are as shown in Fig 5A. The same DosR + AcP data set is shown in panels (B) and (C). p-values were obtained with a 2-way ANOVA with Tukey’s multiple comparisons. Significant p-values are shown for the PknH-DosR + AcP, or PknD-DosR + AcP, versus DosR + AcP proteins. ** p<0.01. The numerical data underlying the graphs shown in this figure are provided in S1 Data.

*dosR* expression is itself upregulated by the very signals that the DosRS(T) system responds to [4, 6, 7]. We found that concentrations as low as 0.25 µM AcP-treated DosR resulted in an increase in target gene transcription rate, with maximal rates obtained at 0.5-1 µM DosR, before levels again decreased at concentrations ≥1.5 µM (Fig 5B, black dashed line). Interestingly, when DosR was phosphorylated by PknH or PknD, in addition to being treated with AcP, an increase in transcription rate was also observed, but with a narrower activation window, as the “turn-down” response occurred with a steeper decline (for example, 4.76 ± 1.67 AU/s and 6.10 ± 1.56 AU/s for 3 µM PknH and PknD-phosphorylated, AcP-treated DosR, respectively, versus 12.14 ± 1.26 AU/s for only AcP-treated DosR, p < 0.01 in each case; Figs 5B and 5C, compare blue and red dashed lines to black dashed line).

These results further demonstrate how STPK phosphorylation can alter DosR activity output, and also intriguingly show a concentration-dependent effect of AcP treatment (HK phosphotransfer) on DosR activity that is modulated by STPK phosphorylation. Together, these results illuminate how STPK phosphorylation of DosR affects its function, and suggests a mechanism by which the response of Mtb to key environmental signals can be tightly regulated.

### STPK phosphorylation sites within DosR affect Mtb transcription and survival in response to environmental signals

Due to the observed mechanistic impact of STPK phosphorylation on DosR function, we next sought to characterize the role of STPK phosphorylation of DosR on Mtb biology. To do this, we first exploited the use of a STPK-phosphoablative DosR variant to examine its impact on Mtb transcriptional response to NO. We generated a STPK-phosphoablative variant of DosR by mutating the T198 and T205 sites to alanine (DosR-T198A/T205A) and performed qRT-PCR analysis on Mtb grown in 7H9, pH 7.0 media ± 100 μM DETA NONOate, an NO donor. Complementation of Δ*dosR* Mtb with the STPK-phosphoablative *dosR* mutant allele (DosR-T198A/T205A) resulted in significantly decreased expression of DosR regulon genes upon NO exposure, compared to complementation with a WT *dosR* allele (Fig 6A).

**Fig 6.**
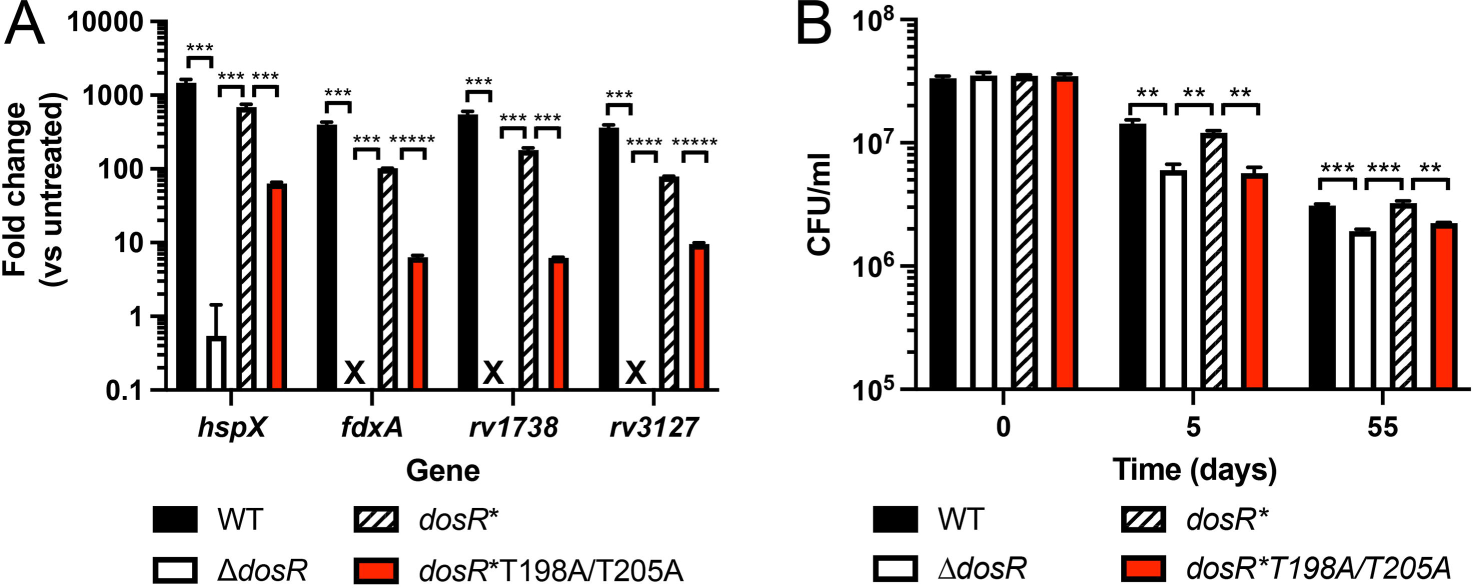
STPK phosphorylation sites within DosR affect Mtb transcription and survival in response to environmental signals. (A) STPK phosphorylation sites within DosR are important for induction of DosR regulon genes in response to NO. Log-phase WT, Δ*dosR*, *dosR\** (complemented mutant), and *dosR\**T198A/T205A Mtb were exposed for 4 hours to 7H9, pH 7 media ± 100 µM DETA-NONOate, before RNA was extracted for qRT-PCR analysis. Fold change compares the 100 µM DETA-NONOate condition to the control untreated condition for each strain. “X” indicates where values for Δ*dosR* were all around -1, and thus not possible to plot on the log-scale. Data are shown as means ± SEM from 3 experiments, and p-values were obtained with unpaired t-tests. *** p<0.001, **** p<0.0001. (B) STPK phosphorylation sites within DosR are important for growth in oxygen-deprived environments. Log-phase WT, Δ*dosR*, *dosR\**, and *dosR\**T198A/T205A Mtb cultures were adjusted to an OD_600_ = 0.3 in fresh 7H9, pH 7 media + 10% oxyrase in sealed tubes and incubated at 37°C. Tubes were taken for colony forming unit (CFU) plating at indicated days post-assay start. Data are shown as means ± SEM from 3 experiments, and p-values were obtained with unpaired t-tests. ** p<0.01, *** p<0.001. The numerical data underlying the graphs shown in this figure are provided in S1 Data.

To test the functional importance of STPK phosphorylation of DosR in a relevant environmental context, we utilized an anaerobiosis assay in which cultures of Mtb were placed into sealed tubes supplemented with oxyrase, which depletes the environment of oxygen [55, 56]. Within 5 days post-assay start, Δ*dosR* Mtb was significantly decreased in its ability to survive in these hypoxic conditions compared to WT Mtb or the complemented *dosR* strain (Fig 6B). Strikingly, complementation of Δ*dosR* with the DosR-T198A/T205A variant failed to rescue the mutant phenotype (Fig 6B).

These results were surprising given our findings above of decreased *hspX* and *fdxA* promoter binding by STPK-phosphorylated DosR (Figs 3C and 3D, Fig 4), and the decreased steady-state transcription observed with *fdxA* (Fig 5). We thus tested the ability of DosR-T198A/T205A to bind DNA and found that this mutant DosR had decreased target DNA binding affinity compared to WT DosR (compare Fig 7A to Fig 3A). Notably, the T198 and T205 sites are located on the l110 helix of each DosR monomer and are closely positioned in the DosR dimer interface in the DNA binding conformation [57] – a proximity that would be predicted to destabilize this mode of dimerization and inhibit DNA binding. Indeed, we further observed that *fdxA* steady-state transcription in the presence of DosR T198A/T205A was comparable to that of PknH-phosphorylated DosR (Fig 7B). Together, these data suggest that either phosphorylation or mutation of these sites inhibit DNA binding due to their unique position in the DosR structure.

**Fig 7.**
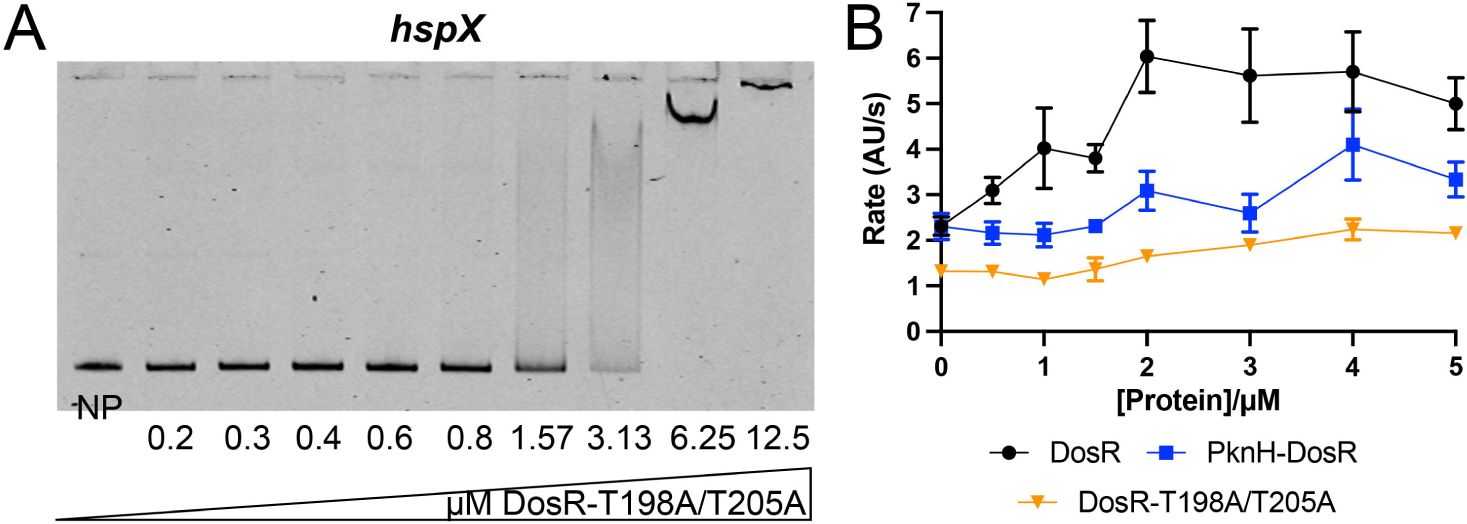
Preservation of DosR STPK phosphorylation sites is important for DosR binding and transcription. (A) EMSA using purified recombinant C-terminally 6x-His-tagged DosR-T198A/T205A and IRDye 700-labeled probes for the *hspX* promoter is shown. A control with no protein (“NP”) added is also shown. DosR-T198A/T205A was added at indicated concentrations for all other lanes. 40 fmoles of *hspX* promoter DNA was used in each reaction. Data are representative of 3 independent experiments. (B) A Spinach RNA aptamer assay was run with the *fdxA* promoter with different concentrations of indicated DosR protein. The WT DosR and PknH-phosphorylated DosR (“PknH-DosR) data are as shown in Fig 5A. Fluorescence (arbitrary units, “A.U.”) was tracked over time on a plate reader, and steady-state rate calculated. Data are shown as means ± SEM from 2-8 experiments. The numerical data underlying the graph shown in this figure are provided in S1 Data.

### The STPK PknH regulates Mtb sensitivity to NO

Given our working hypothesis, supported by the observations above, that STPK activity is functionally relevant to regulation of the Mtb stress response, we finally sought to examine how *pknH* deletion might alter the sensitivity of Mtb to NO. To this end, we transformed WT, Δ*pknH*, and *pknH\** (complemented mutant) with our NO/hypoxia-responsive *hspX’*::GFP reporter [5], and tested reporter response in different DETA-NONOate concentrations. Markedly, *hspX’*::GFP reporter induction was higher at an intermediate DETA-NONOate concentration (50 µM) with Δ*pknH* Mtb as compared to WT or *pknH*\* Mtb (Fig 8). In contrast, in the presence of 100 µM DETA NONOate, reporter induction was similar across all strains (Fig 8).

**Fig 8.**
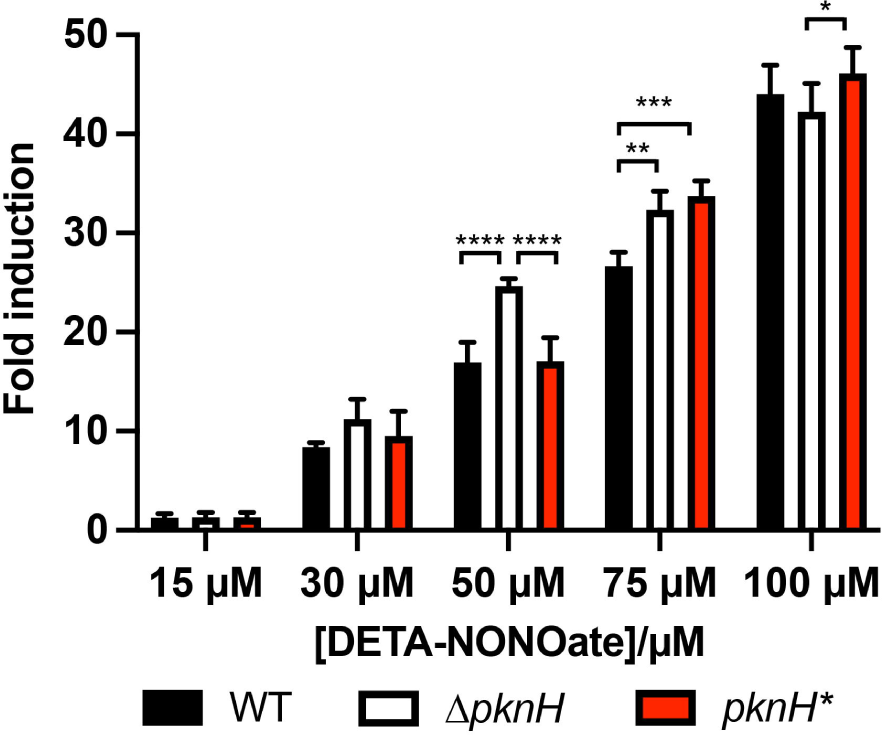
The STPK PknH regulates Mtb sensitivity to NO. WT, Δ*pknH*, and *pknH*\* (complemented mutant) Mtb each carrying the *hspX’*::GFP reporter were exposed for 1 day to the indicated DETA-NONOate concentrations, before fixation and reporter signal analysis via flow cytometry. Fold induction is in comparison to the untreated condition for each strain. Data are shown as means ± SEM from 3 experiments. p-values were obtained with a 2-way ANOVA with Tukey’s multiple comparisons. Only significant comparisons are indicated. * p<0.05, ** p<0.01, *** p<0.001, **** p<0.0001. The numerical data underlying the graph shown in this figure are provided in S1 Data.

These results again support a role of PknH as a fine-tuning regulatory mechanism for Mtb environmental response, acting to restrain DosR activation to ensure response initiation only when appropriate.

## DISCUSSION

The ability of Mtb to sense and respond to environmental cues is dependent on signal transduction regulatory mechanisms such as STPKs and TCSs, which are critical for bacterial adaptation and hence survival within a host [14, 21, 22, 24, 27, 58–60]. While the interplay between STPKs and TCSs has become increasingly appreciated, how such interplay may alter TCS function and thus the response of Mtb to environmental cues has remained understudied. Here, we uncover a role of STPK phosphorylation of TCSs as a fine-tuning regulatory mechanism that acts to provide an additional layer of regulation of TCS RR activity.

In particular, our finding that PknH phosphorylation of DosR decreases DNA binding affinity to target promoters and results in decreased steady-state transcription of target genes strongly supports that regulation of DosR by STPKs serve as a second axis of regulation, in addition to regulation by its cognate histidine kinases (HKs). A previous study had conversely reported increased binding of PknH-phosphorylated DosR to the *hspX* promoter [36], but differed from our assays in testing binding to separate DosR boxes present in the *hspX* promoter (38 bp or 54 bp segments), versus the longer DNA segments encompassing all DosR boxes upstream of the *hspX* promoter utilized in our study. Notably, both the DosR T198 and T205 sites phosphorylated by PknH map within the α10 helix of the DosR crystal structure [61, 62]. DosR is thought to adopt two dimeric structures, one active and one inactive, that exist in equilibrium with each other as well as a monomeric form [57]. Phosphotransfer to the D54 site in the receiver domain favors the active, DNA-binding DosR, with the α10 helix from each monomer mediating the dimerization of that species, making this helix critical for protein activation [57, 61]. We posit that PknH phosphorylation at sites located within this helix can shift the equilibrium of DosR in such a way as to disfavor the active DNA binding-competent dimer form, resulting in the decreased DNA binding affinity and steady-state transcription observed. Examples of opposing effects of STPK and HK phosphorylation on activity of the corresponding RR have been previously reported in *S. aureus* and *Streptococcus agalactiae* [41, 42], and our data further support this concept of STPKs serving to restrain TCS activity to ensure appropriate response.

Additionally, our results with the DosR-T198A/T205A variant indicate the importance of the location of STPK phosphorylation sites on TCS RRs, and indeed, known or putative STPK phosphorylation sites for the Mtb TCS RRs MtrA and PhoP are also positioned close to or within the DNA binding region [32, 60, 63, 64]. In the case of the Mtb TCS RR PrrA, STPK phosphorylation occurs within the receiver domain, affecting Mtb response to various environmental signals [24]. Such positioning of STPK phosphorylation sites close to or within the DNA binding or receiver regulator domains are also found in other bacterial species [37, 38, 42]. Future studies seeking to uncover how STPK phosphorylation alters TCS structure and hence function will be important for continued insight into the role of STPKs as regulators of TCSs.

A key outstanding question is how STPK activity is regulated. Expression levels of STPKs have not generally been found to be affected by key environmental signals for Mtb, and Mtb encodes only a single Ser/Thr phosphatase, with broad activity across phosphorylated Ser/Thr residues [27]. The large number of targets for a given STPK would however strongly suggest a need for regulation of their activity, and indeed the basal activity level for many STPKs appears low, as evidenced by deletion mutants of the STPKs exhibiting little reduction in phosphosites, compared to WT Mtb, in standard rich media [32]. STPKs autophosphorylate and dimerization is required for their activation [27, 65, 66]; ligand triggering of such events thus represent a route for control of STPK activity separate from expression differences. Indeed in *B. subtilis*, the STPK PrkC senses cell wall fragments, with binding of peptidoglycan fragments to its extracellular domain leading to phosphorylation and activation of an essential ribosomal GTPase involved in initiating vegetative growth [67]. In Mtb, a Δ*pknG* mutant is defective for growth when glutamate or asparagine is used as the sole nitrogen source, and phosphorylation of GarA, a key target of PknG, was greatest in the presence of these same amino acids, suggesting that these amino acids may act as triggers for PknG activity [68]. Intriguingly, we found almost complete overlap in the STPKs that interact with the PrrA and DosR RRs, both known to sense and respond to NO and hypoxia [24, 25, 69]. A smaller subset of those same STPKs also interacted with PhoP, which like PrrA, functions as a global regulator of pH and Cl^-^ [5, 24].

Future studies analyzing the extent of phosphorylation of each RR by a given interacting STPK identified here, and testing whether the same environmental signals that drive TCS activity also affect STPK activity, will provide important insight into the coordination of TCS and STPK activity.

Finally, the RNA aptamer-based transcriptional assay utilized here provides a real time, quantitative, and high throughput method with broad utility for understanding basal and regulated transcription dynamics. This encompasses the ability to test effects of different intrabacterial signals and regulatory factors on gene transcription kinetics, through to analysis of antibiotic-dependent inhibition of RNA polymerase on transcription steady-state rates [53, 54]. Interestingly, this transcriptional assay revealed a non-monotonic relationship between AcP-activated DosR concentration and target gene steady-state transcription rates. More specifically, after the expected increase in transcription with initial increases in DosR concentration, we observed an unexpected decrease in transcription with further DosR concentration increases. In addition, the slope of this decrease appeared to be modulated by STPK phosphorylation. In Mtb, expression of the DosR regulon is markedly elevated upon initial exposure to NO or hypoxia [4, 6, 7], reflecting its role in mediating the early transcriptional response to these stresses. However, this induction is transient, and expression levels decline over time even in the continued presence of the inducing signal [70]. Our results suggest a possible negative feedback mechanism incorporating DosR phosphorylation state, whereby active DosR concentrations above a threshold result in decreased target gene expression relative to the maximum activation at lower concentrations. The HK DosS has been reported to also possess phosphatase activity [71, 72]; defining how such phosphatase activity may work in concert with both the observed RR concentration dependence and STPK phosphorylation to enable a negative feedback loop in DosR regulon expression will be vital for understanding physiological adaptation of Mtb to environmental cues that are maintained over the course of a chronic infection.

TCSs and STPKs are the two major regulatory systems through which environmental signals are transduced into adaptive outputs in bacteria. Our findings here illustrate how STPK-mediated phosphorylation of TCS RRs can act to fine tune transcriptional outputs, serving to ensure response initiation only when appropriate. Our work establishes a framework for dissecting STPK-TCS interplay, and we propose that further studies probing the interactions of these two regulatory systems will continue to yield important insight into molecular pathways critical for Mtb environmental adaptation and host colonization.

## MATERIALS AND METHODS

### Mycobacterial protein fragment complementation assays

Mycobacterial protein fragment complementation (M-PFC) assays were performed essentially as described previously [46]. The open reading frames of *dosR*, *prrA*, *phoP*, and *kdpE* were each cloned into pUAB100, generating C-terminal translational fusions to the murine dihydrofolate reductase (mDHFR) fragment F1,2 domain. All Mtb STPK genes were cloned into pUAB400, generating N-terminal translational fusions to the mDHFR fragment F3 domain. For STPKs with transmembrane domains (all except PknG and PknK), only the kinase domain was cloned. *M. smegmatis* transformed with each respective plasmid pair (each STPK with each RR) were plated on 7H11 plates supplemented with 25 µg/ml kanamycin and 50 µg/ml hygromycin, with TRIM added at 10, 20, 30, or 50 µg/ml. pUAB100 and pUAB400 plasmids containing the *Saccharomyces cerevisiae* GCN4 homodimerization domain served as the positive control [46]. As a negative control, *M. smegmatis* expressing the RR-mDHFR F1,2 fusion in pUAB100 with an empty pUAB400 plasmid was used. *M. smegmatis* transformants carrying mDHFR F3-PknB or mDHFR F3-PknI could not be obtained, and M-PFC assays with these STPKs were thus not pursued.

### Recombinant protein expression and purification

The open reading frames of DosR and DosR-T198A/T205A were individually cloned into the isopropyl-β-D-1-thiogalactopyranoside (IPTG)-inducible pET-23a vector to generate a C-terminally 6xHis-tagged DosR and DosR-T198A/T205A, respectively. Expression constructs for the kinase domains of PknH and PknD were previously described [66]. All constructs were transformed into *Escherichia coli* BL21(DE3) for recombinant expression and purification. For expression, 2 ml of overnight *E. coli* cultures started from single colonies were grown in 5 ml LB + 50 μg/ml ampicillin at 37°C and were used to inoculate 1 L LB media + 50 μg/ml ampicillin. Cultures were grown at 37°C, 160 rpm, until the culture reached an OD_600_ of ∼0.6. Induction of constructs was initiated by adding 1 mM IPTG, and the cultures grown for an additional 16 hours at 16°C, 160 rpm. Afterwards, supernatants were removed, and pellets were stored at -80°C prior to further processing.

Recombinant purification of DosR and its variants and STPKs followed previously described protocols [1, 24]. To remove phosphorylated residues accrued from expression in *E. coli*, the protein was treated with alkaline phosphatase (Sigma #P0114) according to previously described protocols [73], when the protein was still bound to the nickel beads. Dephosphorylated DosR was then washed three times with a minimal low imidazole buffer (500 mM NaCl, 50 mM Tris, pH 7.5, 15 mM imidazole, 10% glycerol) to remove residual alkaline phosphatase. For *in vitro* phosphorylation of DosR, nickel bead-bound DosR was treated with 1 µM PknH or PknD and incubated at room temperature in kinase buffer (40 mM Tris-HCl, pH 7.5, 2 mM MnCl_2_, 20 mM MgCl_2_, 2 mM DTT, 0.5 mg/mL BSA) for one hour prior to continuation of the protein purification protocol. DosR protein was dialyzed into electrophoretic mobility shift assay (EMSA) buffer as described [45]. Protein concentrations were quantified by using a Bradford assay (Bio-Rad).

Mtb RNAP σ^A^ holoenzyme complex was purified by a 10X N-terminal His-tag on the alpha subunit, using pET-Duet-*rpoB*-*rpoC,* pAcYc-His*rpoA*-*rpoZ*, and pAC27-*sigA* plasmids, expressed and purified as previously described [54]. The final holoenzyme fractions were dialyzed into storage buffer (10 mM Tris-Cl, pH 7.0, 200 mM NaCl, 0.1 mM EDTA, 1 mM MgCl_2_, 20 μM ZnCl_2_, 2 mM DTT, 50% glycerol), concentrated to 4.5 μM (determined using an extinction coefficient of 280,425 M^-1^ cm^-1^), aliquoted, flash frozen in liquid nitrogen, and stored at -80°C.

Mtb CarD and RbpA, in pET-SUMO plasmid vectors, were expressed, purified, and the His-SUMO tag removed as previously described [54, 74]. Eluted fractions were dialyzed overnight in 20 mM Tris, pH 8.0, 150 mM NaCl, 1 mM β-mercaptoethanol, then concentrated to 200 μM determined using extinction coefficients of 16,900 M^-1^ cm^-1^ for Mtb CarD and 13,980 M^-1^ cm^-1^ for Mtb RbpA.

### Mass spectrometry

Protein samples underwent in-gel trypsin digestion at the Mass Spectrometry Technology Access Center at the McDonnell Genome Institute (MTAC@MGI) at Washington University School of Medicine. The peptides were subject to mass spectrometry to confirm the phosphorylation on DosR without phosphopeptide enrichment. The sample data was searched for phosphorylation on D/S/T/Y residues against a custom *E. coli* BL21 (DE3) database plus the sequence of DosR. In all samples, DosR accounted for at least 80% of the identified peptides and 99% sequence coverage of DosR was obtained in all samples.

### Electrophoretic mobility shift assays

Electrophoretic mobility shift assays were performed essentially as previously described [1, 75]. In brief, promoter regions for *hspX* (558 bp) and *fdxA* (221 bp) were amplified using IRDye 700 labeled primers (Integrated DNA technologies) and the PCR products purified using a QIAquick PCR purification kit (Qiagen). For acetyl phosphate-treated reactions, purified DosR-6xHis protein was treated as described previously [45]. Indicated amounts of unphosphorylated and phosphorylated DosR were mixed with 40 fmoles of DNA in EMSA buffer (25 mM Tris-HCl, pH 8, 20 mM KCl, 6 mM MgCl_2_, 5% glycerol, 0.5 mM EDTA, 25 µg/ml salmon sperm DNA [45]) in a 10 µl final reaction volume. Reactions were incubated at 20°C for 30 minutes and then run on non-denaturing 7% Tris-glycine gels in 0.5X Tris-borate-EDTA buffer at 4°C for 4 hours. Gels were imaged using the 700 nm channel of an Odyssey CLx imaging system (LI-COR).

### Fluorescence polarization assays

Fluorescence polarization (FP) experiments were performed using linear double-stranded DNA probes containing either the *hspX* (85 bp) or *fdxA* (55 bp) promoter labeled with Alexa Fluor 488 on the downstream 5’ end (Integrated DNA Technologies). These were titrated with increasing concentrations of purified DosR, or PknH/PknD-treated DosR. 10 µl binding reactions in 384-well black, low volume, round bottom assay plates (Corning) were sealed with optical adhesive film (Applied Biosystems) and measured on a CLARIOstar Plus plate reader (BMG LABTECH). DosR and its phosphorylation variants were serially diluted (1:1.5) from a 25 µM stock solution into binding buffer (25 mM Tris-HCl 8.0, 20 mM KCl, 6 mM MgCl_2_, 10% glycerol), generating a final concentration range of 0-25 µM. Labeled DNA substrates were added to each well at a final concentration of 25 nM (5 µl of 50 nM stock). Plates were incubated at 37°C for 15 minutes before polarization measurements were taken using 485/520 nm excitation/emission FP filters.

For analysis, binding curves were fit to a Hill-type saturation equation using maximum likelihood estimation (MLE) implemented in R. Raw data were plotted, and fits were generated using the following Hill equation: ΔFP = (ΔFP_max_⋅[P]^n^)/ (K_d_ ^n^ + [P]^n^); Where ΔFP is the change in fluorescence polarization, [P] is the protein concentration (µM), ΔFP_max_ is the maximal signal, K_d_ is the apparent dissociation constant, and n is the Hill coefficient. For the DosR condition, where the data reach saturation, all three parameters were allowed to float. For the PknH-DosR and PknD-DosR conditions, which did not reach saturation, ΔFPmax was fixed to the value obtained from DosR alone, and MLE was used to estimate K_d_ and n, and the residual variance error parameter, σ. 95% confidence intervals (CIs) for parameter estimates were obtained by bootstrapping. For each condition, the Hill model was refit to 10000 synthetic datasets generated by data resampling. The 2.5^th^ and 97.5^th^ percentiles of the resulting distributions of parameter estimates defined the CI limits for each parameter. These confidence intervals were propagated to the fitted curves, producing shaded ribbons in the plots that visualize uncertainty in the predicted fluorescence polarization across the protein concentration range. On the *hspX* promoter the residual variances (σ) were as follows: 16.3 ± 0.98 (DosR), 5.52 ± 0.47 (PknH-DosR), and 13.2 ± 0.91 (PknD-DosR). On the *fdxA* promoter, σ values were: 12.1

± 0.89 (DosR), 12.9 ± 1.27 (PknH-DosR), and 10.0 ± 0.54 (PknD-DosR).

*hspX* probe:

/5Alex488N/ACAACAGGGTCAATGGTCCCCAAGTGGATCACCGACGGGCGCGGACAAATGGCCCGCGCTTCGGGGACTTCTGTCCCTAGCCCTG

*fdxA* probe:

/5Alex488N/TGACGAATAAGGCCTTTGGTCCTTTCCGGTAGGGGTCTTTG GATAGGCGCGATCC

### RNA aptamer-based transcriptional assay

Aptamer-based transcription data was collected using a CLARIOstar Plus Microplate reader (BMG LABTECH) in a 384 well, low volume, round-bottom, non-binding polystyrene assay plate (Corning) with the corresponding Voyager analysis software and following a previously published protocol [54]. To measure multi-round, steady-state transcription kinetics in real-time, we monitored the change in 3,5-difluoro-4-hydroxybenzylidene imidazolinone (DFHBI) fluorescence upon binding to a transcribed, full-length RNA sequence containing the *fdxA* promoter and the iSpinach D5 aptamer. DFHBI fluorescence was measured using an excitation wavelength of 480 ± 15 nm (monochromator) while monitoring the emission signal at 530 ± 20 nm (filter). All reactions were conducted at 37°C in 10 μl final volume in 20 mM Tris (pH 8.0 at 37°C), 40 mM NaCl, 75 mM potassium glutamate, 10 mM MgCl_2_, 5 μM ZnCl_2_, 20 μM EDTA, 5% glycerol with 1 mM DTT and 0.1 mg/ml BSA. Reactions contained 100 nM RNAP holoenzyme, 20 μM DFHBI dye (Sigma Aldrich), 0.4 U/μl RiboLock RNase inhibitors (Thermo Scientific), CarD and RbpA at 1 μM and 2 μM, respectively, and dilutions of DosR from 1 μM to 50 μM. 2.5 μl stock rNTPs (Thermo Scientific) were injected *in situ* using the reader’s automated reagent injector to a final concentration of 1 mM NTP. Data were acquired in 10-20 second intervals for up to 40 minutes total. A minimum of 3 technical replicates of the negative control (*i.e.*, no rNTPs) were collected and measured concurrently with the experimental data. Using the average of this negative control, the experimental data was corrected as previously described [53], bringing all starting fluorescence values to zero and correcting for any time-dependent drift in fluorescence. Between 2 and 8 independent experiments were collected for each condition with 3 technical replicates each. Standard deviations were used as a statistical weight during the linear regression analyses as previously described to obtain the steady-state rate [53].

### Mtb strains and culture

Mtb CDC1551 was used as the parental strain for all assays here, and Mtb cultures were cultured and maintained as described previously, with 7H9 broth supplemented with 10% OADC, 2% glycerol, 0.05% Tween-80, and 100 mM MOPS used for buffering to pH 7.0 [76]. Generation of Δ*dosR*, Δ*pknH*, and their complements were constructed with methods as described previously [5]. The *dosR* deletion consisted of a region from the beginning of the open reading frame through nucleotide 650, while the *pknH* deletion encompassed the entire *pknH* open reading frame. Complementation in both cases utilized the respective endogenous promoters and open reading frames in the integrating plasmid pMV306. The *dosR\**T198A/T205A mutant was generated using QuikChange mutagenesis (Agilent). The *hspX’*::GFP reporter introduced into indicated strains was previously reported [5]. Antibiotics were added as needed at the following concentrations: 100 μg/ml streptomycin, 50 μg/ml hygromycin, 50 μg/ml apramycin, and 25 μg/ml kanamycin.

### qRT-PCR analyses

Mtb grown to log-phase (OD_600_ ∼ 0.6) in aerated conditions was used to inoculate filter-capped T75 flasks laid flat, containing 12 ml 7H9, pH 7.0 ± 100 μM DETA NONOate at OD_600_ = 0.3. Bacteria were incubated for 4 hours, before RNA extracted as previously described [3]. qRT-PCR experiments were conducted and analyzed according to previously established protocols [77].

### Anaerobiosis assay

Mtb strains were propagated to mid-log phase (OD_600_ ∼ 0.6) and subcultured to OD_600_ = 3 in a 2 ml screw cap tube with 1.1 ml 7H9, pH 7.0 media. 0.13 ml of oxyrase was added to reduce oxygen. [56], and tubes were tightly capped, wrapped with parafilm, and incubated at 37°C. Each tube was used only for a single time point, with serial dilutions plated on 7H10 agar plates containing 100 μg/ml cycloheximide for colony forming unit quantification.

### *hspX’*::GFP reporter assay

Indicated Mtb strains carrying the *hspX’*::GFP reporter were propagated to log phase (OD_600_ ∼ 0.6) and subcultured to an OD_600_ = 0.05 in flat T75 flasks with filter caps containing 4 ml 7H9, pH 7. After 4 passages, Mtb was subcultured at OD_600_ = 0.05 in 7H9, pH 7.0 media with 0, 15, 30, 75, or 100 μM DETA NONOate (Cayman Chemicals). 1-day post-exposure, culture aliquots were taken and fixed in 4% paraformaldehyde. Reporter signal was analyzed via flow cytometry as previously described [77].

### Statistical analyses

GraphPad Prism software was used for all statistical analyses, with p < 0.05 considered significant. The statistical test used for a given assay is described in the figure legends.

## ACKNOWLEDGEMENTS

We thank Christopher Sassetti (University of Massachusetts Medical School) for generously providing STPK expression plasmids, and Adrie Steyn (University of Alabama at Birmingham and Africa Health Research Institute) for the M-PFC plasmids. We thank Elizabeth Billings and Janessa Ya for assistance with the M-PFC assays. We thank the Mass Spectrometry Technology Access Center at the McDonnell Genome Institute (MTAC@MGI) at Washington University School of Medicine for mass spectrometry analyses.

## SUPPORTING INFORMATION FIGURE LEGEND

**S1 Fig. Changes in DosR phosphorylation status alter DNA binding affinity to the promoter of its target gene *fdxA*.** Electrophoretic mobility shift assays (EMSAs) using purified recombinant C-terminally 6x-His-tagged DosR and IRDye 700-labeled probes for the *fdxA* promoter are shown. A control with no protein (“NP”) added is shown for each gel. DosR was added at indicated concentrations for all other lanes. 40 fmoles of *fdxA* promoter DNA was used in each reaction. EMSAs shown are as follows: (A) untreated DosR, (B) DosR incubated with 50 mM acetyl phosphate (AcP), (C) DosR phosphorylated “on-bead” with 1 µM PknH, (D) DosR phosphorylated “on-bead” with 1 µM PknD, (E) DosR phosphorylated “on-bead” with 1 µM PknH, then purified and incubated with 50 mM AcP, and (F) DosR phosphorylated “on-bead” with 1 µM PknD, then purified and incubated with 50 mM AcP. Data are representative of 3 independent experiments.

**S1 Data. Numerical data underlying the presented graphs.** Excel file with numerical data underlying graphed average data presented.

