## Supplementary figures and images for "Serine/threonine protein kinase phosphorylation of DosR alters target gene transcription mechanics and regulates *Mycobacterium tuberculosis* sensitivity to nitric oxide stress"

### S1 Fig

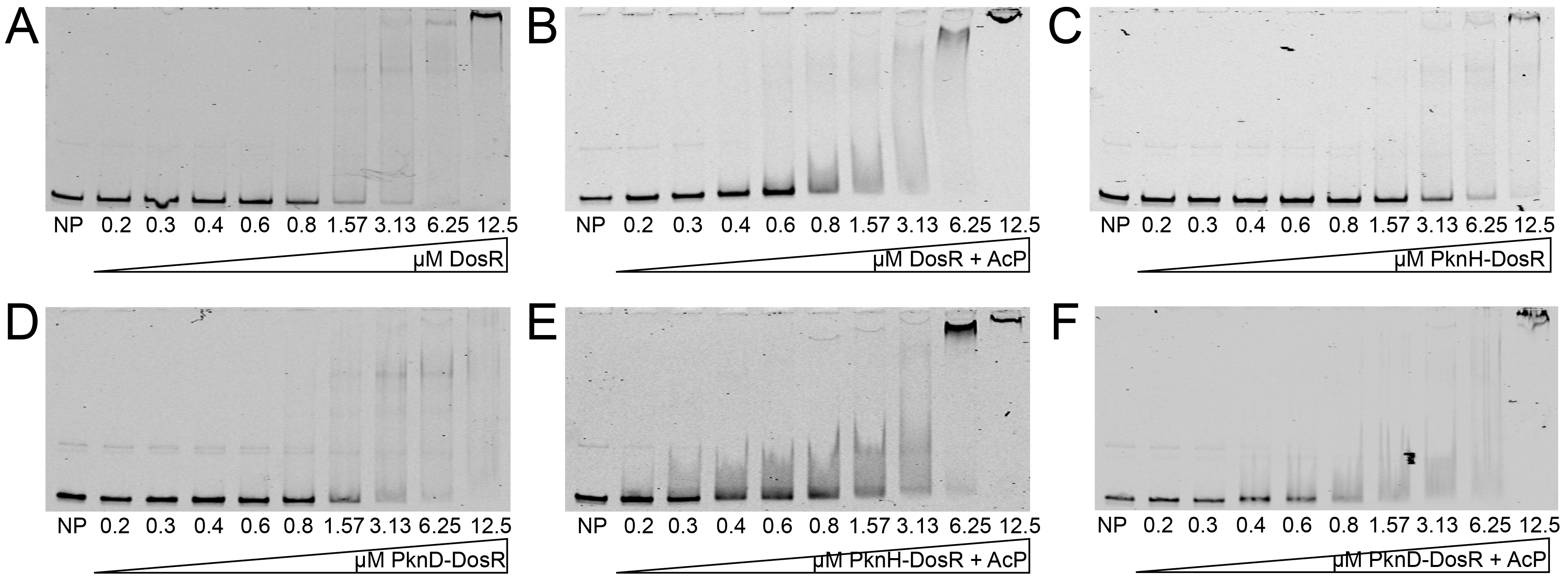
